# RSAD2/VIPERIN and CMPK2 Coordinate an Immunometabolic Response to Epstein-Barr Virus

**DOI:** 10.64898/2026.02.04.703699

**Authors:** Urvi S. Zankharia, Adam M. Glass, Qing Zhu, Janvhi Suresh Machhar, Ying Ye, Bhanu Chandra Karisetty, Jayamanna Wickramasinghe, Andrew Kossenkov, Jozef Madzo, Sun Sook Chung, Samantha S. Soldan, R. Jason Lamontagne, Lawrence Harris, Tyler L Grove, Steven Jacobson, Chengyu Liang, Paul Lieberman

## Abstract

Epstein-Barr Virus (EBV) infection and reactivation in B-lymphocytes is tightly regulated by host antiviral response genes. In the present study, we identify interferon stimulated genes RSAD2 (radical S-adenosyl methionine domain-containing 2) and CMPK2 (Cytidine/Uridine Monophosphate Kinase 2) as key modulators of EBV expression and cellular response during EBV infection and reactivation. EBV primary infection and reactivation lead to a coordinated up-regulation of RSAD2 and CMPK2. Depletion of RSAD2 reduced cell viability and limited EBV reactivation, while depletion of CMPK2 led to reactivation of EBV lytic gene expression during latency. Transcriptomic analysis revealed that RSAD2 and CMPK2 have overlapping functions in regulating IFN-signaling pathways, as well as oxidative phosphorylation, protein translation, and unfolded protein response during reactivation. Despite distinct subcellular localizations, RSAD2 at the Endoplasmic Reticulum (ER), and CMPK2 in mitochondria, both genes converge on shared immunometabolic pathways, including control of Gasdermin D (GSDMD) associated pyroptosis and ATF-4 associated unfolded protein response (UPR). EBV reactivation induced formation of antiviral ribonucleotide ddhCTP during lytic EBV reactivation which was strictly dependent on RSAD2. Knockdown of RSAD2 and CMPK2 had significant effects on global metabolites consistent with a remodeling of glycolysis, fatty acid biosynthesis and degradation of superoxides. These observations demonstrate that RSAD2-CMPK2 function in a coordinated ER-mitochondria stress-Interferon signaling axis that shapes EBV reactivation and host immune control, including a novel layer of immunometabolic regulation modulating viral latency and reactivation.

**Authors Summary:** Understanding how Epstein-Barr virus (EBV) regulates host factors to control infection, latency and reactivation is critical for developing targeted therapies against EBV-associated diseases. This study identifies Interferon Stimulated Genes RSAD2 (Viperin) and CMPK2 as key regulators of EBV reactivation and host interferon responses in B-cells. Despite distinct organelle localizations, both genes converge on a shared immunometabolic pathways, revealing a coordinated ER-mitochondria axis that shapes viral expression and immune signaling. These findings provide new insights into the roles of host antiviral effectors and uncover potential targets for modulating EBV activity in inflammatory and oncogenic contexts.

## Introduction

Epstein–Barr virus (EBV), a member of the γ-herpesvirus family, persistently infects more than 90% of the global population and is implicated in a range of malignancies including B-cell lymphomas, nasopharyngeal carcinoma and gastric carcinoma, collectively accounting for over 200,000 cases of cancer annually [1, 2]. Beyond oncogenesis, EBV infection has been strongly associated with autoimmune disorders such as Multiple Sclerosis (MS), where it is considered an essential cofactor [3]. A critical yet incompletely understood aspect of EBV biology is the role of interferon (IFN) signaling and interferon-stimulated genes (ISGs) in regulating viral latency, lytic reactivation, and host immune response. IFN-signaling induces a broad repertoire of ISGs that establish an antiviral state by restricting viral replication at multiple stages of the viral life cycle [4]. How IFN antiviral response controls EBV infection and may contribute to latency establishment is not completely understood.

RSAD2 (encoding Viperin, Virus Inhibitory Protein Endoplasmic Reticulum Associated Interferon-Inducible) and CMPK2 (Cytidine/Uridine Monophosphate Kinase 2) are two ISGs located immediately adjacent to each other on chromosome 2p25 and cotranscribed during IFN-signaling [5]. Both are expressed as a fusion protein in some lower organisms [6]. RSAD2/Viperin is highly evolutionarily conserved radical S-adenosylmethionine (SAM) enzyme with broad-spectrum antiviral activity. RSAD2/Viperin catalyzes the conversion of cytidine triphosphate (CTP) into 3′-deoxy-3′,4′-didehydro-CTP (ddhCTP), a nucleotide analog that acts as a chain terminator for viral RNA-dependent RNA polymerases, thereby inhibiting replication of viruses such as Zika virus, West Nile Virus and SARS-CoV2 [6–8]. Beyond its antiviral role, RSAD2 is increasingly recognized as a biomarker in autoimmune diseases, including Systemic Lupus Erythematosus (SLE) [9], Rheumatoid Arthritis [10] and in various cancers, where its expression correlates with poor prognosis [10, 11]. RSAD2 is highly expressed in mouse dendritic cells (mDCs) and is necessary for IRF-7 mediated maturation of mDCs [12]. Depletion of RSAD2 lead to loss of mDCs’ ability to induce proinflammatory cytokine production and T-cell proliferation in pulmonary metastasis [12]. CMPK2, a mitochondrial nucleotide kinase, catalyzes the phosphorylation of cytidine and uridine monophosphates to their diphosphate forms, providing essential precursors for mitochondrial DNA synthesis [1, 5]. CMPK2 is crucial for maintenance of macrophage homeostasis and polarization [13]. Overexpression of CMPK2 promotes microglial activation and neuroinflammation through cGAS-STING pathway, implicating it in neurodegenerative disease mechanisms [14].

Despite these insights, the roles of RSAD2 and CMPK2 in EBV infection and host gene regulation remain poorly understood. We show here that RSAD2 and CMPK2 are coordinately regulated during EBV infection and reactivation in B-lymphocytes. To elucidate their function in EBV biology, we employed lentiviral-mediated knockdown of RSAD2 and CMPK2 in EBV+ Burkitt Lymphoma (BL) cell lines to assess their impact on viral gene expression and host immune response. We found that RSAD2 and CMPK2 orchestrate distinct yet overlapping roles in regulating EBV reactivation and host innate immune responses. Their knockdown disrupts interferon response, upregulates pathways and metabolites involved in mitochondrial metabolism and alters unfolded protein response dynamics, highlighting a coordinated ER-mitochondria-immune axis that shapes EBV pathogenesis and host defense.

## Results

### RSAD2 and CMPK2 are upregulated during EBV primary infection, lytic reactivation, and in MS-patient derived SLCLs with unstable latency

Previous studies from our lab examined EBV infection of primary B-cells isolated from whole blood of normal healthy donors at various days after infection (**Fig 1A**) [15]. Further analysis of the published RNAseq data revealed significant activation of RSAD2 and CMPK2 transcripts at days 2, 7 and 21 post-infection (**Fig 1B**). RSAD2 and CMPK2 are known ISGs that share a common genetic locus on chr 2p25. Examination of chromatin features revealed a strong ATAC-seq activation of a single peak situated in the common promoter/enhancer regulatory region of the two divergently transcribed genes (**Fig. 1C**). ChIP-seq peaks for EBNA1, CTCF, and H2A.Z overlapped with the ATAC-seq peak suggesting that EBNA1 may directly regulate the chromatin accessibility and transcription of RSAD2 and CMPK2 (**Fig. 1C**). We confirmed that EBNA1 bound to the RSAD2/CMPK2 promoter/enhancer locus by ChIP-qPCR (**Fig. 1D**). Thus, EBV infection strongly induces expression of both RSAD2 and CMPK2 in primary B-cells through a mechanism involving EBNA1 binding and increased chromatin accessibility in the common promoter/enhancer regulatory region.

**Figure 1:**
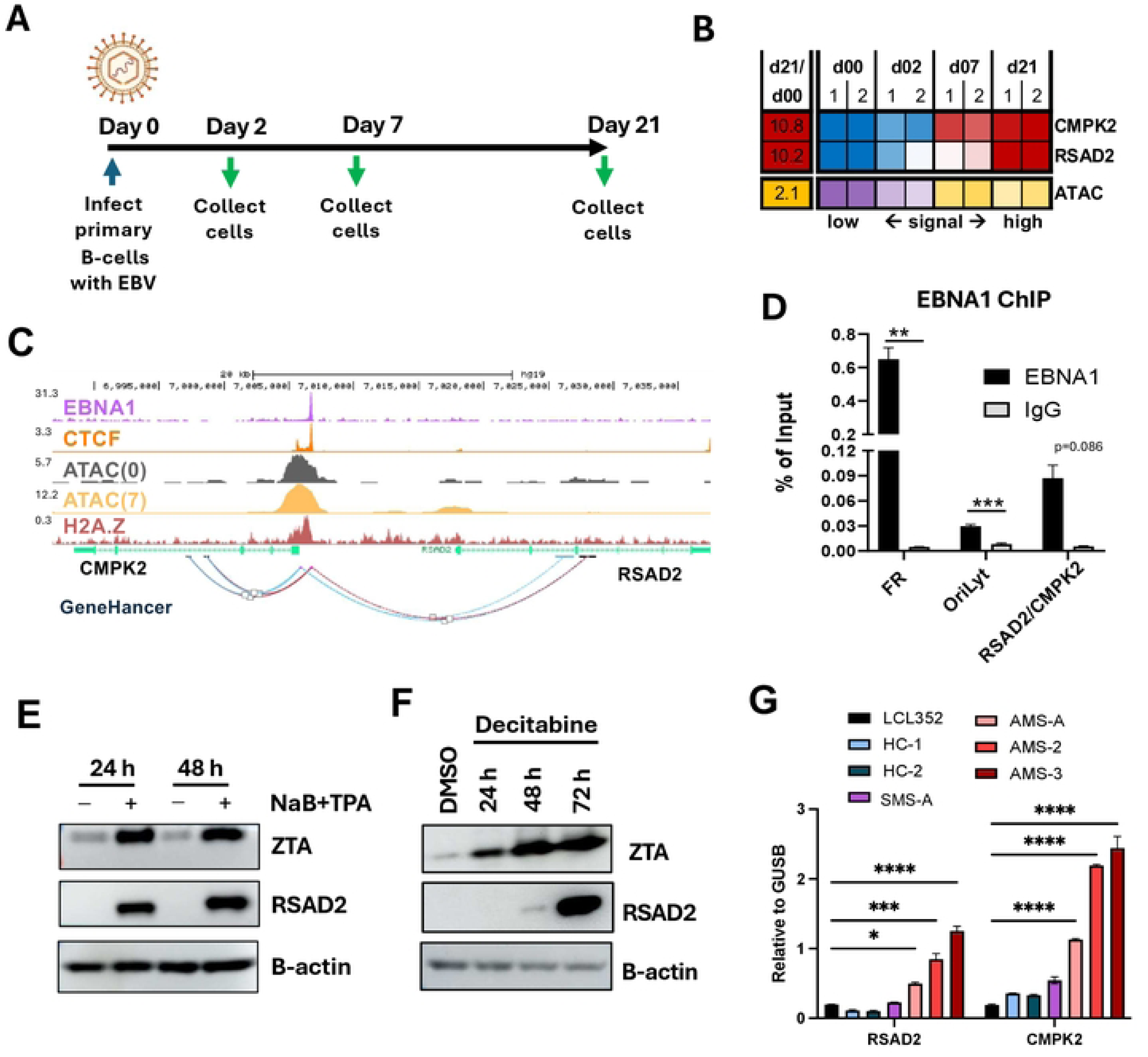
RSAD2 and CMPK2 are upregulated during EBV primary infection, lytic reactivation, and in Multiple Sclerosis (MS)-patient derived spontaneous lymphoblastoid cell lines (SLCLs) with unstable latency. **A)** Timeline of EBV infection (Mutu I derived) in primary B-cells. **B)** RNA-seq and ATAC- seq signals after EBV infection of primary B-cells. **C)** Track representation with gene model and genehancer tracks in the context of EBNA1, CTCF, and H2A.Z ChIP-Seq and ATAC-Seq signal at day 0 and 7 post EBV infection. **D)** ChIP-qPCR assay for EBNA1 binding to EBNA1/CTCF binding site at the RSAD2/CMPK2 locus. **E-F)** Western blot analysis in Mutu I after treatment with sodium butyrate (NaB) and TPA (E) or Decitabine (F). **G)** Expression analysis by RT-qPCR for RSAD2 (left) or CMPK2 (right) relative to GUSB in LCL352 or SLCLs derived from MS patients with active disease (AMS), stable disease (SMS) or healthy controls (HC). Statistical analysis was performed using ordinary two-way ANOVA using Sidak’s multiple comparisons (panels D and G). n=3, * p<.05, ** p<.01, ***p<.001, ****p<.0001.

We next examined RSAD2 protein expression during latency and reactivation in EBV+ BL cells. RSAD2 protein was undetectable in latently infected Mutu I cells (**Fig 1E**), but readily detected after induction with NaB+TPA (**Fig 1E**) or by Decitabine (**Fig. 1F**) as measured by Western blot. This demonstrates that RSAD2 is also upregulated during viral reactivation from latency.

We next tested whether RSAD2 and CMPK2 expression correlates with the intrinsic control of EBV latency. A previous study found that spontaneous lymphoblastoid cell lines (SLCLs) derived from MS patients had unstable latent infections with increased lytic cycle gene expression and inflammatory cytokine production compared to healthy controls [16]. We examined this cohort of SLCLs and found that patients with active MS (AMS) had significantly higher expression of both RSAD2 and CMPK2 as compared to LCL352 (Mutu I transformed) and SLCLs derived from healthy controls (HC) or stable disease (SMS) **(Fig 1G**). Expression levels of both genes were up to 10-fold higher in patients with active disease (**Fig 1G**) where lytic gene expression and inflammatory cytokines are elevated [16]. These findings suggest that RSAD2 and CMPK2 expression correlate with the EBV lytic activity and inflammatory state associated with active MS.

### Knockdown of RSAD2 modulates expression of EBV

To investigate the effect of RSAD2 on EBV and cellular response during latency and lytic reactivation, we knocked down RSAD2 in Mutu I. We used three different shRNAs in either uninduced latency or after treatment with NaB+TPA to induce lytic EBV reactivation (**Fig 2A)**. RSAD2 was efficiently knocked down by all three shRNAs as measured by Western blot in Mutu I cells induced by NaB+TPA (**Fig. 2B**). More sensitive RT-qPCR analysis of RSAD2 transcript also demonstrated efficient knockdown of RSAD2 in uninduced Mutu I cells (**Fig 2C**). Knockdown of RSAD2 decreased cell viability relative to control shRNA by ∼50%, suggesting that RSAD2 is essential for Mutu I cell survival (**Fig 2D**). Thus, although present in exceedingly low amounts under uninduced latency, RSAD2 is essential for cell survival. RSAD2 knockdown in latently infected Mutu I cells led to a modest increase in expression of EBV transcripts (**Fig 2E**). Induction of EBV reactivation by treatment with NaB+TPA upregulated the expression of RSAD2 protein and transcript ∼100-fold as compared to uninduced cells (**Fig 2F compared to 2C**), and shRNA depletion was highly efficient, with virtually no detectable RSAD2 protein (**Fig. 2B**) or RNA (**Fig. 2G**). While RSAD2 is thought to provide antiviral function, we found that knockdown of RSAD2 during lytic induction significantly reduced EBV lytic transcription, reducing expression of ZTA and EA-D by >80% (**Fig 2G**). This suggests that RSAD2 is required for efficient EBV transcription during lytic induction and that loss of it severely compromises EBV expression.

**Figure 2:**
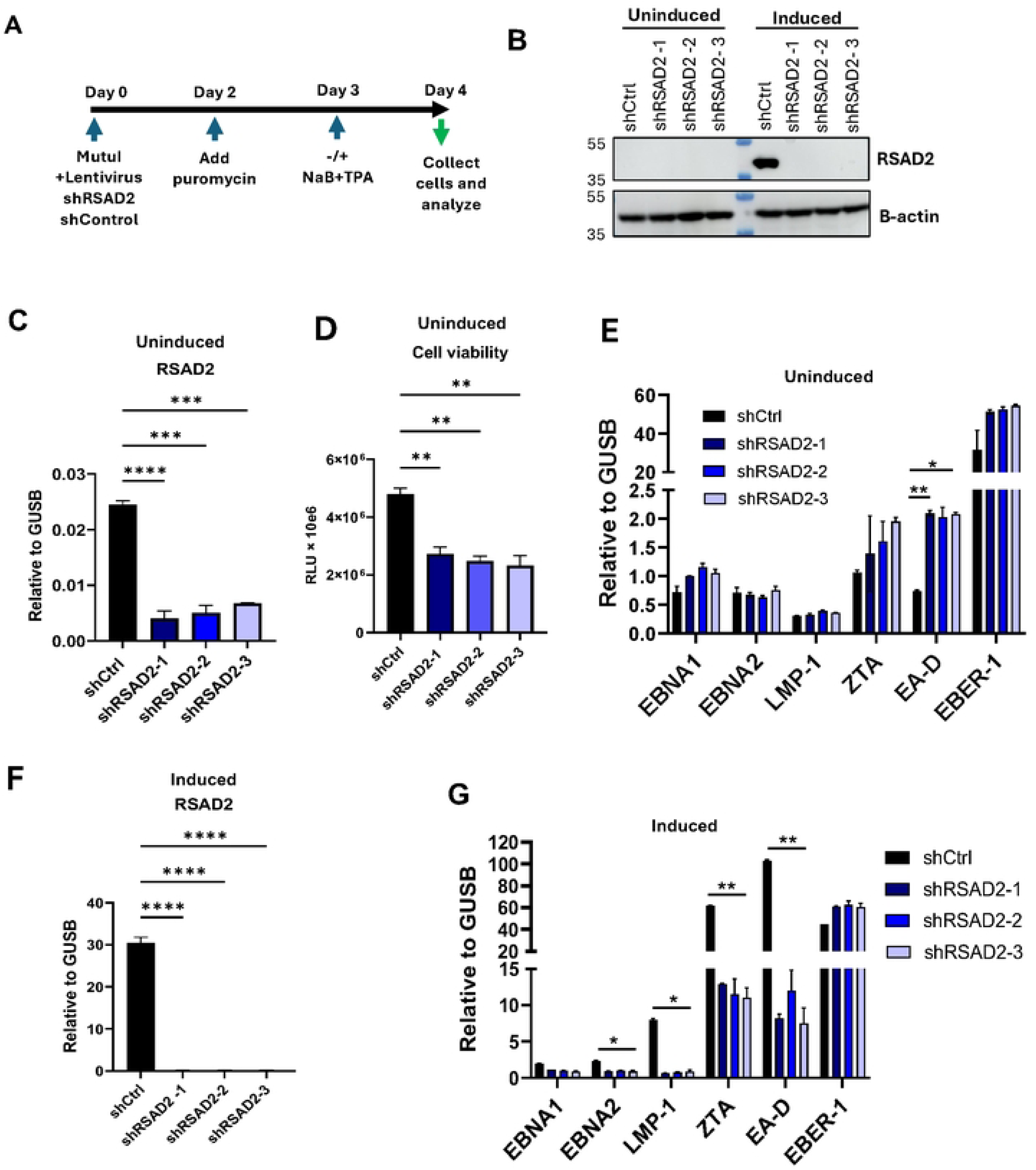
Knockdown of RSAD2 modulates EBV reactivation and cell viability. **A**) Schematic representation of timeline for RSAD2 knockdown by lentivirus transduction in Mutu I. **B)** Western blot for RSAD2 expression in uninduced and induced Mutu I with shCtrl or three independent shRNA for shRSAD2. **C**) RT-qPCR analysis for RSAD2 expression after knockdown in uninduced Mutu I. **D**) Measurement of cell viability by CellTiterGlo assay in uninduced Mutu I after RSAD2 knockdown. **E**) EBV expression analysis by RT-qPCR in uninduced Mutu I. **F**) RT-qPCR analysis for RSAD2 expression after knockdown in induced Mutu I. **G**) RT-qPCR for EBV mRNA expression in induced Mutu I. Statistical analysis was performed using ordinary one-way ANOVA using Dunnett’s multiple comparisons (panels C,D and F) or ordinary two-way ANOVA using Sidak’s multiple comparisons (panels E and G). Data shown here is representative of results from three independent experiments. n=3, * p<.05, ** p<.01, ***p<.001, ****p<.0001.

### CMPK2 is antiviral and restricts EBV expression

To study the effect of CMPK2 on EBV expression and cellular response, we knocked down CMPK2 using three different shRNAs in Mutu I using experimental timeline as shown in **Fig 3A**. Under basal conditions, expression of CMPK2 is low and all three shRNAs efficiently knocked down CMPK2 (**Fig 3B**). Knockdown of CMPK2 did not have significant impact on cell survival (**Fig 3C**). Depletion of CMPK2 led to significant upregulation of EBV lytic gene expression in uninduced Mutu I cells as shown by RT-qPCR (**Fig 3D**), as well as a significant increase (∼30 fold) in EBV DNA copy number indicating lytic replication was occurring (**Fig. 3E**). Western blot also revealed an increase in lytic cycle protein ZTA and EA-D, as well as latency proteins EBNA1 and LMP1 (**Fig. 3F**). Knockdown of CMPK2 during lytic EBV reactivation reduced CMPK2 mRNA transcripts by ∼50% (**Fig. 3G**), and led to minor changes to EBV transcripts, including increases in EBNA1, EBNA2, and EBER1, and decrease in LMP1 as detected by RT-qPCR (**Fig. 3H**). Interestingly, Western blot showed an increase in EA-D and LMP1 protein, along with an increase in RSAD2/Viperion protein, consistent with the enhanced reactivation of EBV and potential translational control of LMP1 and RSAD2/Viperin (**Fig. 3I**). These findings suggest that CMPK2 acts as a restriction factor for EBV during latent infection in Mutu I, and modulator of LMP1 and RSAD2/Viperin protein translation during lytic reactivation.

**Figure 3:**
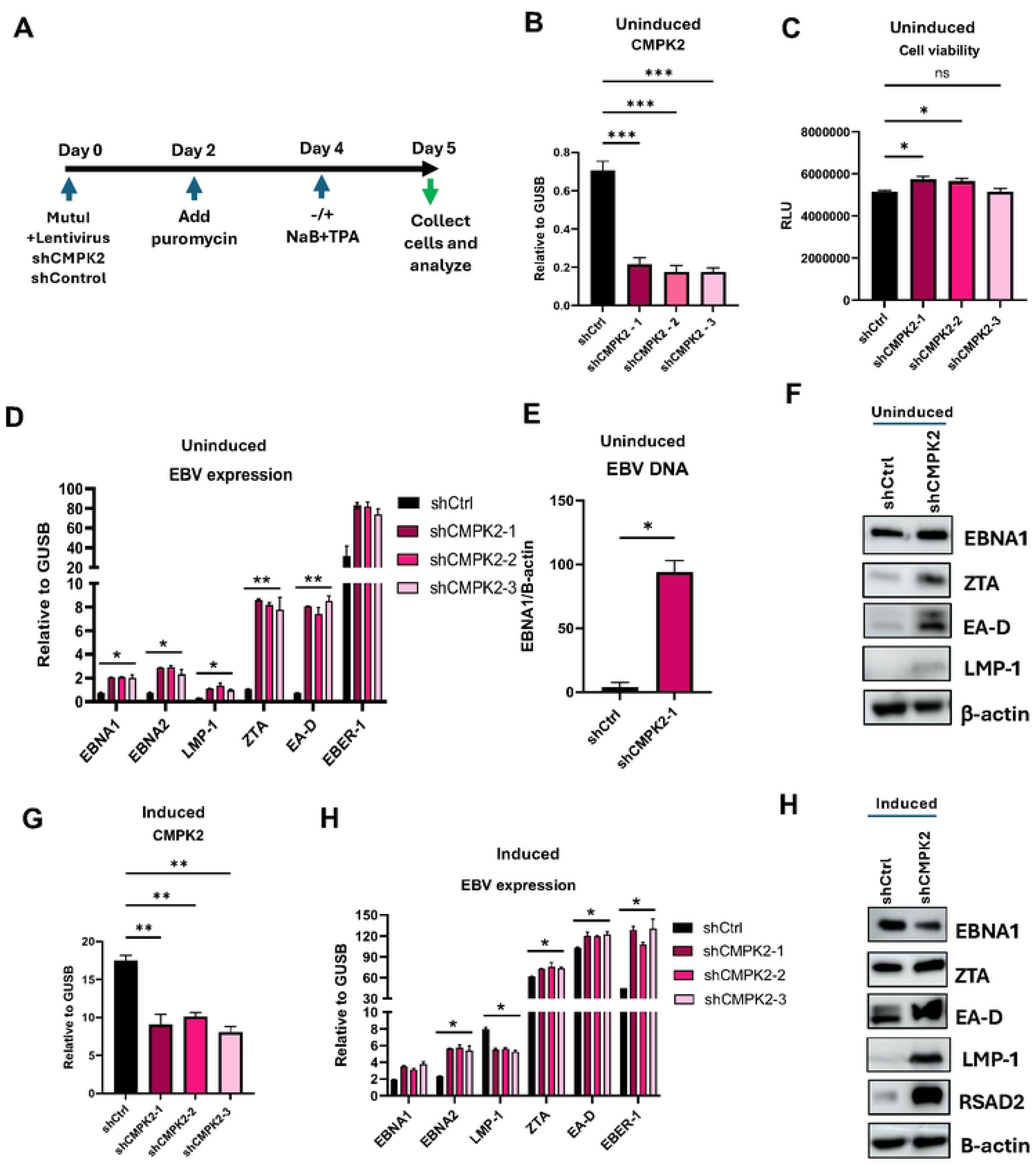
CMPK2 is antiviral and restricts EBV lytic expression. **A**) Schematic representation of timeline for CMPK2 knockdown in Mutu I. **B**) RT-qPCR analysis for CMPK2 expression after knockdown in uninduced Mutu I. **C**) Measurement of cell viability by CellTiterGlo assay in uninduced Mutu I. **D**) EBV mRNA expression analysis by RT-qPCR in uninduced Mutu I. **E**) EBV DNA copy number measured by qPCR for EBNA1 gene relative to β-actin. **F**) Western blot analysis for EBV protein expression for uninduced Mutu I. **G**) RT-qPCR for CMPK2 expression after knockdown for induced Mutu I. **H**) EBV mRNA expression analysis by RT-qPCR for induced Mutu I. **I**) Western blot analysis for EBV protein expression for induced Mutu I. Statistical analysis was performed using ordinary one-way ANOVA using Dunnett’s multiple comparisons or ordinary two-way ANOVA using Sidak’s multiple comparisons. Data shown here is representative of results from three independent experiments n=3, * p<.05, ** p<.01, ***p<.001, ****p<.0001.

### Knockdown of RSAD2 leads to apoptosis while depletion of CMPK2 enhances cell survival

Since we found that RSAD2 knockdown led to substantial cell death, we further wanted to study the effect of RSAD2 and CMPK2 knockdown on apoptosis. We analyzed for apoptosis by flow cytometry using the Annexin V/7-AAD staining (**Fig. 4A**). We found that loss of RSAD2 during uninduced and induced conditions lead to substantial cell death (**Fig 4A and B**). Under uninduced conditions, depletion of RSAD2 in Mutu I reduced cell viability from 78% to 39%, which accounts to ∼50% decrease in live cell population (**Fig 4A and B**). Interestingly, while knockdown of CMPK2 under uninduced conditions had no significant effect on cell viability, depletion of CMPK2 during lytic EBV reactivation significantly enhanced cell viability (**Fig 4A and B**). Thus, CMPK2 may enhance EBV reactivation by preventing antiviral apoptotic response.

**Figure 4:**
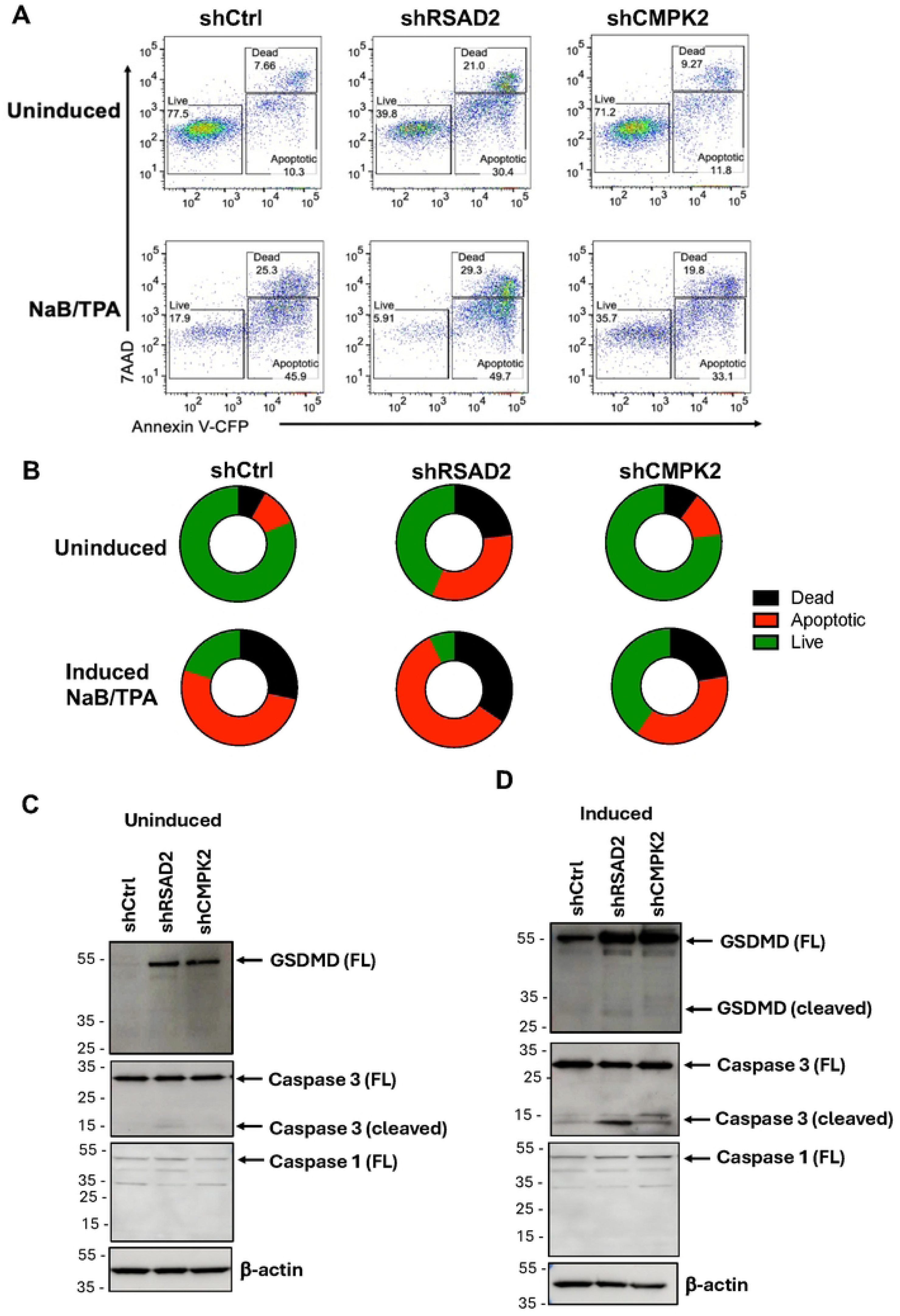
Knockdown of RSAD2 leads to apoptosis while depletion of CMPK2 enhances cell survival. **A**) Apoptosis assay by flow cytometry after Annexin V-7AAD staining in inuced or uninduced Mutu I transduced with shCtrl, shRSAD2, or shCMPK2. **B**) Circle chart of live, dead, and apoptotic cells after treatment shown in panel A. **C-D**) Western blot analysis for expression of GSDMD, Caspase 1, Caspase 3, and loading control β-actin under uninduced condition (C), and induced condition (D).

To better understand the role of RSAD2 or CMPK2 modulation on cell viability, we performed Western blot analyses. We found Gasdermin (GSDMD), a marker for pyroptosis [17], was upregulated after RSAD2 and CMPK2 knockdown during uninduced (**Fig. 4C**) and induced conditions (**Fig. 4D**). Caspase 3 cleavage, a marker of apoptosis and pyroptosis [18], was detected after RSAD2 knockdown in uninduced and a more substantial cleavage signal in induced conditions (**Fig. 4C and D**). Caspase 1 cleavage, a marker of inflammasome activity [19], was not detected under these conditions. Thus, a specialized mechanism of cell death, possibly involving pyroptosis, occurs after knockdown of RSAD2 in reactivating Mutu I cells.

### RSAD2 and CMPK2 are essential for induction of efficient IFN-response during lytic EBV reactivation

To gain additional insight into the mechanism of RSAD2 and CMPK2 in regulating cellular response during latency and lytic EBV reactivation, we performed transcriptional profiling by RNA-seq after knockdown of either RSAD2 or CMPK2 (**Fig. 5 and S1-5**). Initial analysis of differential gene expression between induced and uninduced identified the upregulation of interferon response and immune cytokine signaling, and down regulation of myc and E2F pathways during EBV lytic reactivation (**Fig. S1**). Knockdown of RSAD2 or CMPK2 had significant effects on both latent and lytic induced cells, but most strikingly were the similar effects on the reduction of interferon signaling during lytic induction (**Fig. 5A, B**, **and E, S2 and S3**). During lytic conditions, CXCL10, CCL22, IFIT2, IFI44L were most significantly downregulated by RSAD2 knockdown (**Fig. 5A, S3**), while CCL22, IFI44, IFI44L, RSAD2, and IFIT3 were most significantly downregulated by CMPK2 knockdown (**Fig. 5B**, **S3**). Knockdown of either RSAD2 or CMPK2 during lytic induction strongly induced expression of ARGHAP32 and LRRC4C, which are implicated in neuronal development and function. We found a highly significant overlap (**Fig. 5C**) and correlation (**Fig. 5D**) in genes regulated by RSAD2 or CMPK2 knockdown under induced conditions. Common pathways most significantly down regulated were interferon gamma and alpha/beta signaling, macrophage Th1 activation pathway, mitochondrial dysfunction, antigen presentation and deubiquitination (**Fig. 5E**). Common pathways most upregulated were oxidative phosphorylation, IL-10 signaling, protein translation, and PD-1/PDL1 immunotherapy pathway and SRP-dependent cotranslational targeting (**Fig. 5E**). Regulators most affected were inhibition of IFNG, TNF and IFNA2P. and activation of MAPK1 and immunity related GTPase IRGM (**Fig. 5F**). Pathways unique to CMPK2 knockdown were related to activation of Eukaryotic translation, EIF2 signaling and inactivation of pre-mRNA processing (**Fig. S4A**). Pathways unique to RSAD2 knockdown were related to activation of cell cycle checkpoints, synthesis of DNA, protein ubiquitination and inactivation of eukaryotic translation initiation and response to EIF2AK4 amino acid deficiency (**Fig. S4B**). We experimentally validated the most affected common genes CXCL-10, CCL22, IFI-6 were reduced by shRSAD2 (**Fig. S5A)** and shCMPK2 (**Fig. S5B**), while RUNX1T1 was upregulated by both RSAD2 and CMPK2 knockdown. Thus, knockdown of RSAD2 and CMPK2 have a highly similar effect on the host transcriptional response to EBV lytic reactivation.

**Figure 5:**
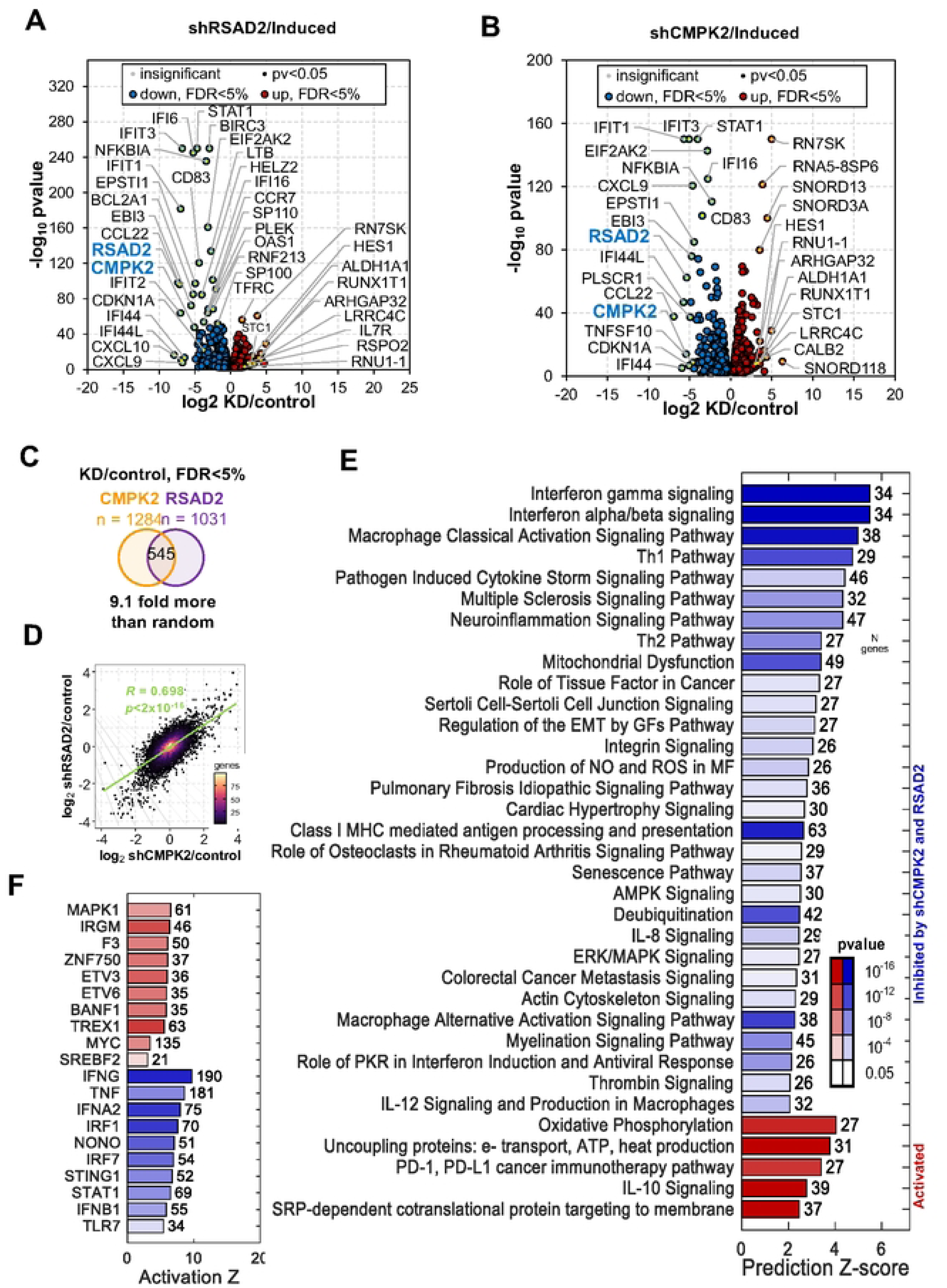
RSAD2 and CMPK2 are essential for induction of efficient IFN-response during lytic EBV reactivation. **A**) Volcano plot showing top significantly changed genes by shRSAD2 in Mutu I induced cells. **B**) Volcano plot showing top significantly changed genes by shCMPK2 in Mutu I induced cells. **C**) Venn diagram showing overlap of commonly regulated genes for shRSAD2 and shCMPK2. **D**) Pearson correlation of transcript changes in shCMPK2 and shRSAD2 in induced Mutu I cells. **E**) Pathways analysis for downregulated (blue) and upregulated (red) genes common to both shCMPK2 and shRSAD2 in Mutu I cells. **F**) Regulators for down (blue) and up (red) activation based on Z-score using IPA.

### Knockdown of RSAD2 and CMPK2 increase mitochondrial transcription and alter ER homeostasis

Since RSAD2 is Endoplasmic Reticulum (ER) localized and CMPK2 is a mitochondrial protein, we wanted to examine the effect of knockdown of RSAD2 and CMPK2 on mitochondrial and ER homeostasis. We first confirmed the intracellular localization of CMPK2 in Mutu I by confocal imaging (**Fig. 6A**). As expected, CMPK2 colocalized with mitochondrial receptor protein Tom 20 [20] and this localization was strongly upregulated during lytic induction (**Fig. 6A** and **B**). Knockdown of CMPK2 upregulated transcription of several mitochondrial genes, especially MT-ND6, MT-CO1 and MT-RNR2 (**Fig. 6C**), suggesting it has a function role in maintaining mitochondrial homeostasis. String database analysis for both CMPK2 and RSAD2 identified each other within their respective networks (**Fig. 6D**). Further, RSAD2 network identified interferon response genes (**Fig. 6D, left**) and CMPK2 network identified many pyrimidine and mitochondrion associated genes (**Fig. 6D, right**). To probe the effects of RSAD2 and CMPK2 knockdown on the interferon signaling pathway, we performed Western blot analyses for known interactors of RSAD2, including IRAK1 and TRAF6, which have been reported as RSAD2 interaction proteins [21]. Knockdown of RSAD2 reduced expression levels of IRAK1, TRAF6 and TAK1 relative to β-actin during induced, and to a lesser extent during uninduced conditions (**Fig. 6E**). Knockdown of CMPK2 increased expression of IRAK1 and TRAF6 under induced conditions as compared to shCtrl (**Fig. 6E**). Since RSAD2 is localized to the ER, and since protein expression was generally decreased after knockdown, we assayed the effects on ER stress and Unfolded Protein Response (UPR) regulatory factors by Western blot analysis (**Fig. 6F**). We found that CHOP and BiP/Grp78, both of which are molecular chaperones localized in the ER, were upregulated after RSAD2 and CMPK2 knockdown in uninduced conditions. Levels of PDI slightly increased after CMPK2 knockdown while ERO1L levels significantly reduced under both uninduced and induced states. ATF-4 levels decreased substantially after lytic induction, and knockdown of either RSAD2 or CMPK2 reduced ATF-4 in uninduced conditions. peIF2 levels were slightly elevated after RSAD2 knockdown in uninduced conditions, further suggesting an effect on protein translation (**Fig 6F**). Thus, knockdown of both RSAD2 and CMPK2 alter the dynamics of ER homeostasis.

**Figure 6:**
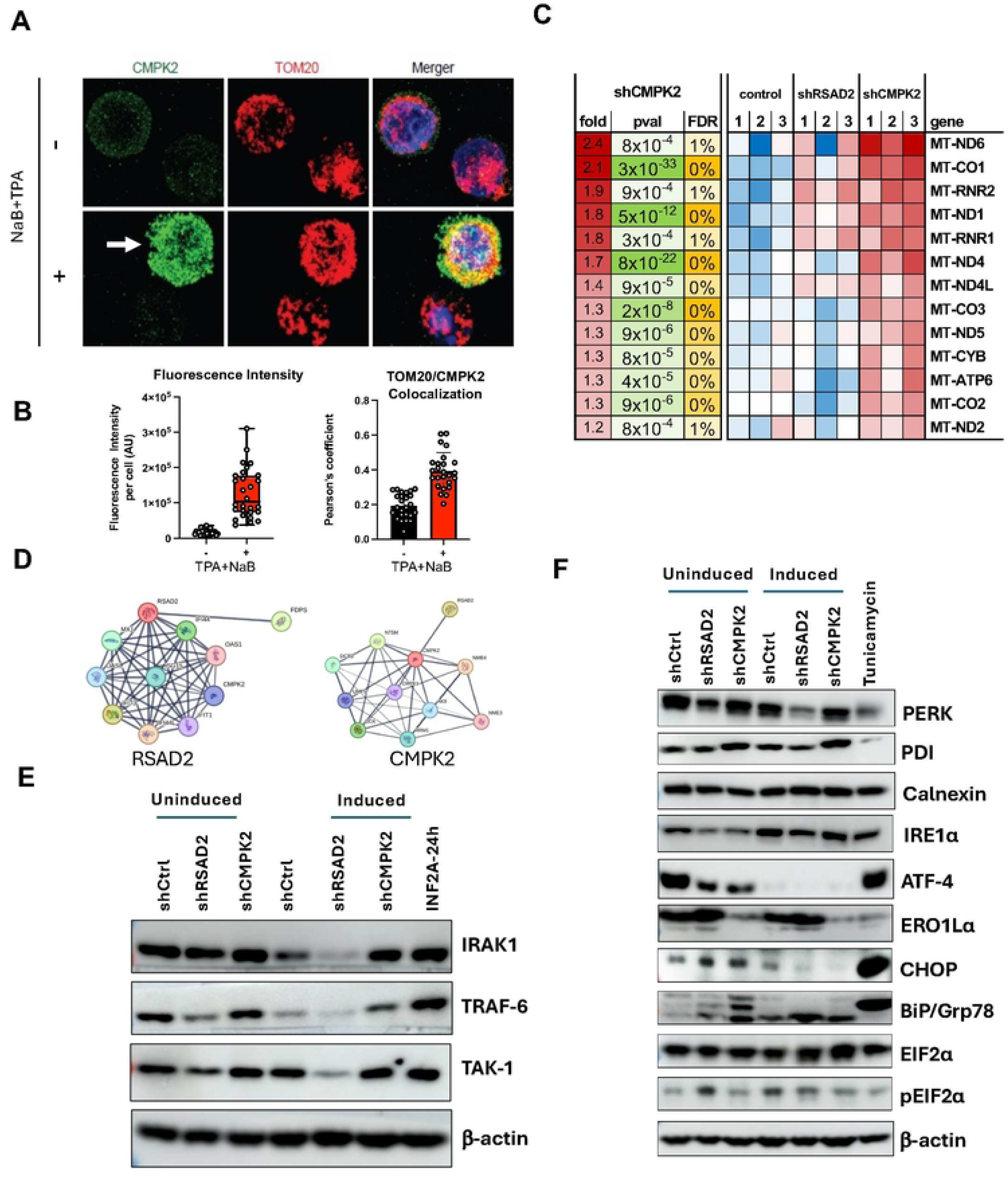
Knockdown of RSAD2 and CMPK2 increase mitochondrial transcription and alter ER homeostasis. **A**) Confocal imaging for CMPK2 and mitochondrial marker Tom20 colocalization with/without NaB+TPA treatment. **B**) Quantification of fluorescence intensity (left) and colocalization events (right) shown in representative image shown in panel A. **C**) Transcriptomic analysis of mitochondrial gene expression by RNA-seq after RSAD2 and CMPK2 knockdown. **D**) String data base interaction map for RSAD2 and CMPK2. **E**) Western blot analysis for expression of interferon signaling proteins IRAK1, TRAF-6, and TAK-1 in uninduced or induced Mutu I cells with shCtrl, shRSAD2, or shCMPK2. **F**) Same as in panel E, but for expression of ER stress markers PERK PDI, Calnexin, IRE1α, ATF-4, ERO1Lα, CHOP, BiP/Grp78, EIF2α, and control β-actin.

### RSAD2-dependent generation of ddhCTP and global metabolite shift during EBV reactivation

To investigate the role of RSAD2 and CMPK2 on metabolic response to EBV, we examined the effects of these enzymes on cellular metabolites. RSAD2 catalyzes the conversion of CTP into ddhCTP, a unique antiviral ribonucleotide that can act as a chain terminator for viral encoded RNA-dependent RNA polymerases [6]. CMPK2 functions to provide CTP substrate for conversion to ddhCTP by RSAD2 [6]. To determine if ddhCTP was produced during lytic EBV reactivation and if knockdown of RSAD2 and CMPK2 affected ddhCTP formation, we used LC-MS to measure ddhCTP levels (**Fig. 7A and B**). LC-MS identification was calibrated using purified synthetic ddhCTP (**Fig. 7A**). Under basal conditions (uninduced), ddhCTP was nearly undetectable (**Fig. 7B**). However, during lytic EBV reactivation, ddhCTP levels were substantially increased in control cells (shCtrl, induced) likely due to increased expression of both RSAD2 and CMPK2 (**Fig. 7B**). Knockdown of RSAD2 during lytic induction led to almost complete loss of ddhCTP, showing that RSAD2 is indispensable for ddhCTP production (shRSAD2, induced). In contrast, knockdown of CMPK2 did not significantly decrease ddhCTP. IFN-2A treated cells showed very high levels of ddhCTP formation, as expected. We also found that EBV reactivation induced cellular levels of ddhCMP (**Fig. S6A**), and ddhCDP (**Fig. S6B**). Interestingly, knockdown of RSAD2 reduced ddhCDP levels (**Fig. S6B**), but not ddhCMP (**Fig. S6A**), suggesting alternative pathways may generate ddhCMP. These findings show the strong correlation between EBV reactivation and the RSAD2 dependent generation of ddhCTP and its related byproducts.

**Figure 7:**
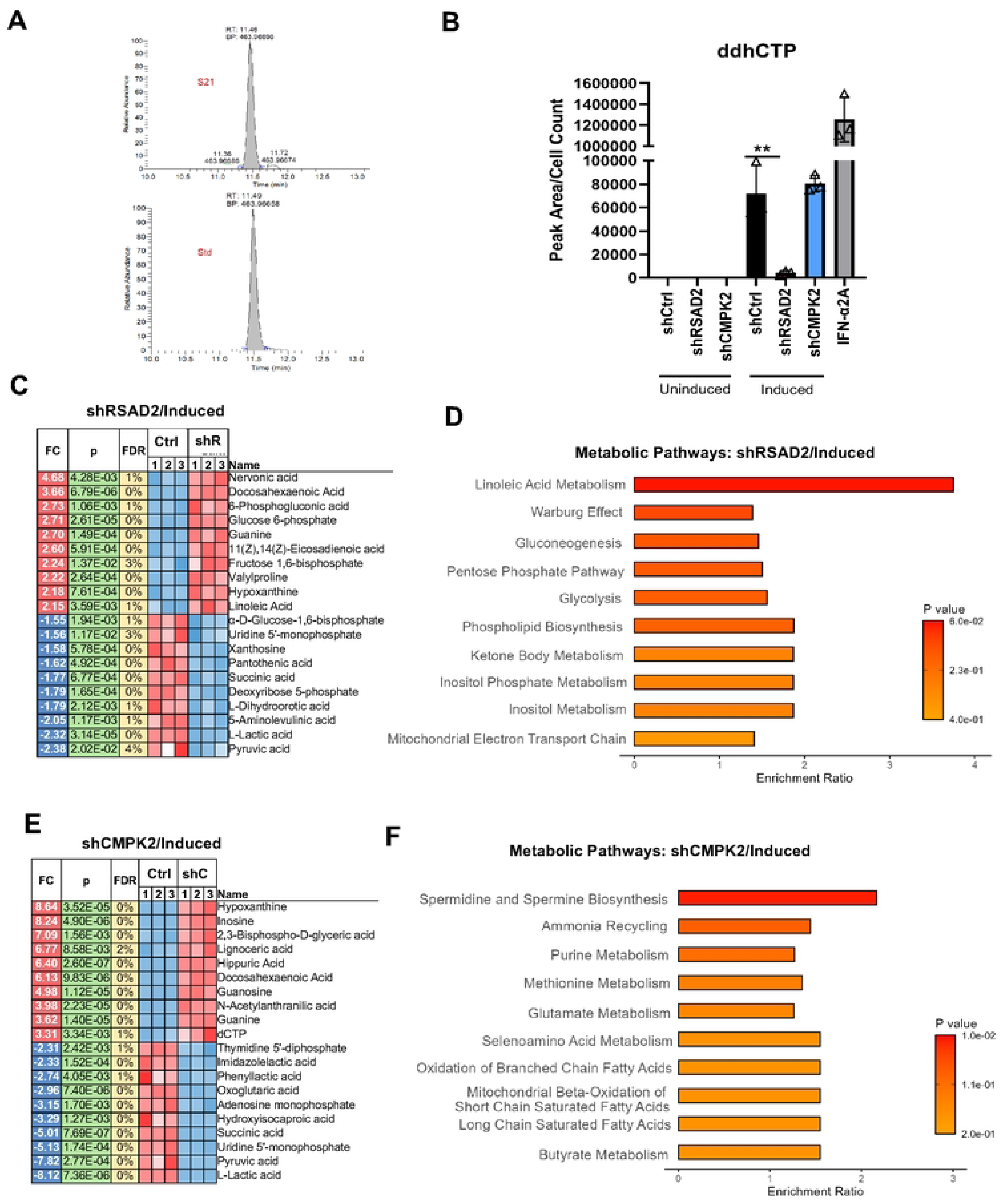
ddhCTP detection after RSAD2 and CMPK2 knockdown and global metabolite shift during EBV reactivation. **A**) LC-MS chromatogram of ddhCTP standard and its detection in the sample. **B**) Detection of ddhCTP in control (shCtrl) RSAD2 knockdown (shRSAD2) and CMPK2 knockdown (shCMPK2) in Mutu I cells under uninduced and induced condition. **C)** Top 10 up/downregulated metabolites after RSAD2 knockdown in induced Mutu I cells. **D**) Pathway enrichment analysis for RSAD2 knockdown in induced Mutu I cells. **E**) Top 10 up/downregulated metabolites after CMPK2 knockdown in induced Mutu I cells. **F**) Pathway enrichment analysis for CMPK2 knockdown in induced Mutu I cells.

To gain a broader understanding of metabolite changes during EBV reactivation and in response to RSAD2 or CMPK2 knockdown, we used mass-spectrometry for global metabolite profiling. Many metabolic changes were observed for each condition (**Fig. 7C-F, S7**). Knockdown of RSAD2 significantly increased levels of Nervonic acid, Docosahexaenoicacid, Linoleic acid, Glucose 6-phosphate, and Fructose 1,6-bisphosphate (**Fig. 7C and D**), while knockdown of CMPK2 increased levels of metabolites such as Hypoxanthine, Inosine, Lignoceric acid, Docosahexaenoicacid, and 2,3-bisphospho glyceric acid (**Fig. 7E and F**). Metabolite changes occurring between uninduced and induced conditions also showed similarities. For example, under uninduced state knockdown of RSAD2 increased levels of Fructose 1,6 bisphosphate, Nervonic acid, and Lignoceric acid (**Fig 7 and S7A**). Metabolites such as Nervonic acid, Lignoceric acid, Docosahexaenoic acid are involved in neurodevelopment, while Glucose 6-phosphate, and Fructose 1-6 bisphosphate are intermediates in glycolysis. Nervonic acid helps in remyelination of nerves in Multiple Sclerosis and also has anti-inflammatory properties. In terms of pathway enrichment, Spermidine and Spermine Biosynthesis had high enrichment ratio for CMPK2 knockdown and, Alphalinoleic acid and Linoleic acid metabolism was enriched in RSAD2 knockdown. There was a significant overlap in the metabolites regulated by both genes including decreased levels of Pyruvic acid, Lactic acid and Succinic acid. We also tested whether interferon treatment altered global metabolites (**Fig. S7E and S7F**). Metabolites that changed most with IFN- treatment included Sepiapterin, Uridine, Cytidine, Lignoceric acid, Docosahexanoic acid, dCMP, dCTP, Lactic acid, Pyruvic acid and Succinic acid. Thus, the metabolic profile overlapped to some extent between IFN--treated cells and knockdown of either RSAD2 and CMPK2, suggesting these enzymes affect common metabolic pathways. These findings demonstrate the RSAD2 and CMPK2 play important roles in regulating cellular metabolism during EBV latency and reactivation in B-lymphocytes.

## Discussion

In the present study, we identified RSAD2/Viperin and CMPK2 as host genes induced by EBV infection and demonstrate their functional roles in modulating host response to viral latency and reactivation. The transcriptional control region of RSAD2 and CMPK2 was bound by EBNA1 and its chromatin accessibility increased by EBV infection. shRNA depletion of RSAD2 and CMPK2 revealed their common function in activation of interferon response genes and maintain host cell survival during EBV reactivation in B-lymphocytes. RSAD2 is induced in EBV infected primary B-cells early during infection (2 dpi). It is possible that at this early and crucial stage of B-cell transformation, RSAD2 might support cell survival and modulate host response to help in efficient expression of early lytic genes and hence facilitate establishment of EBV latency. Depletion of CMPK2 led to a spontaneous reactivation of latent EBV suggesting it functions as an EBV lytic cycle restriction factor. Depletion of RSAD2 led to a loss of viability during EBV reactivation through a mechanism involving cleavage of Caspase 3 and induction of Gasdermin D, suggesting a manner of protection against apoptosis and pyroptosis. RSAD2 is localized to the ER and its depletion led to an activation of ER stress and UPR. CMPK2 localized to the mitochondria and its depletion led to an increase in mitochondrial transcripts. Thus, RSAD2 and CMPK2 function together in the innate immune control of EBV infection and latency (**Fig 8**).

**Figure 8.**
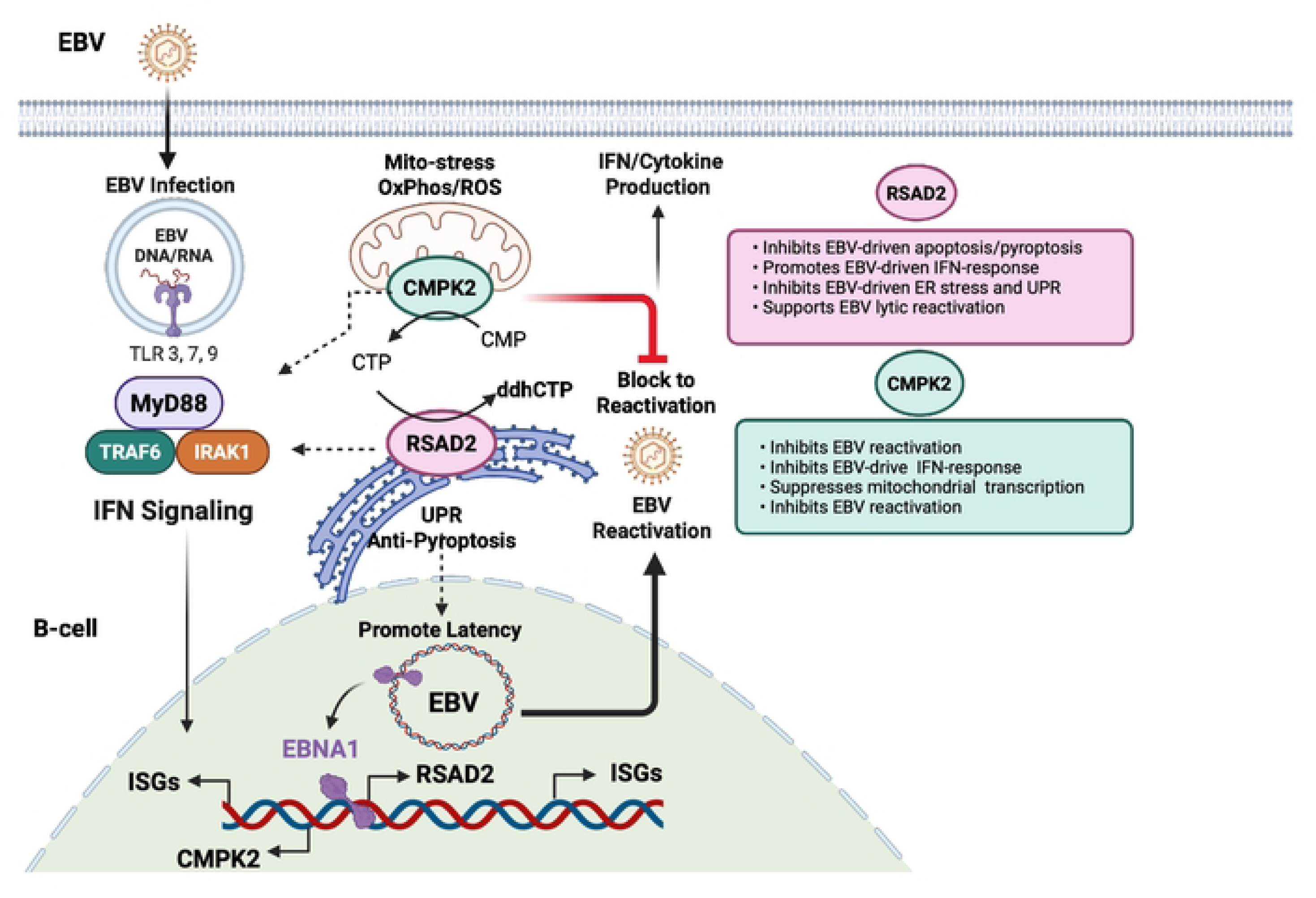
Model showing the interrelated role of RSAD2 and CMPK2 in regulating immune-metabolic response to EBV latency and reactivation.

RSAD2 and CMPK2 are coordinately regulated during EBV infection and lytic reactivation of B-cells. We found that EBNA1 can bind to the common promoter region regulating the divergently transcribed genes, and that ATAC-seq chromatin accessibility increased during EBV infection. EBNA1 binding site overlapped with histone H2A.Z and in close proximity to CTCF suggesting complex and higher order chromatin regulation. RSAD2 and CMPK2 are known IFN-stimulated genes (ISGs) that can be induced directly by interferon signaling. Thus, EBV is likely to directly modulate expression of RSAD2 and CMPK2 through EBNA1 and other viral factors.

Multiple studies have demonstrated that RSAD2/Viperin is highly upregulated following viral infection and has been shown to primarily restrict viral replication [5, 6, 22]. However, other studies, particularly with DNA viruses, have found a proviral function for RSAD2/Viperin. RSAD2/Viperin was found to support Kaposi’s sarcoma-associated herpesvirus lytic replication through methionine oxidation and stabilization of the viral helicase [23]. In HCMV, virally rerouted RSAD2/Viperin relocalizes to mitochondria where it inhibits ATP production, activating AMPK, and triggering lipogenesis to support enveloped virion assembly [24]. Thus, RSAD2/Viperin is usurped for a proviral function during CMV infection. In a previous study conducted by our group in HIV-infected macrophages, we found reduction in HIV transcripts and p24 viral capsid protein levels after RSAD2 knockdown, showing that RSAD2 is essential for sustained HIV infection of human macrophages [25]. In our present study, we found a reduction in EBV lytic transcription after RSAD2 knockdown, partly due to the increase in apoptosis/pyroptosis and activation of UPR. This clearly demonstrates the importance of RSAD2 in promoting EBV lytic gene expression. Thus, RSAD2 is a proviral survival factor for EBV lytic transcription.

In contrast to RSAD2, CMPK2 knockdown increased EBV expression under both latent and lytic conditions, positioning itself as a restriction factor in B-cells. Apoptosis assay revealed that RSAD2 depletion triggers substantial cell death without strong inflammasome activation markers such as NLRP3. We observed accumulation of full-length Gasdermin D and cleaved Caspase 3, which are markers for Pyroptosis/Apoptosis [17]. This reflects the threshold of caspase or inflammasome activation required to induce cell death. In contrast, CMPK2 knockdown increases cell viability during lytic reactivation. Interestingly, we found that knockdown of CMPK2 led to a reduction in RSAD2 transcripts during lytic reactivation in Mutu I, but a substantial increase of RSAD2/Viperin protein. The increase in Viperin protein level after CMPK2 knockdown could reflect post-transcriptional regulation- for example, reduced proteasomal turnover, altered translation selectivity or compartmental retargeting. Increased EBV expression after CMPK2 knockdown may also be the reason for increased RSAD2 protein levels because EBV upregulated RSAD2 expression. Whether increased RSAD2/Viperin protein after CMPK2 knockdown is protective or permissive needs further investigation. Since RSAD2/Viperin is an important survival factor, it is possible that high Viperin protein accumulation after CMPK2 knockdown may be increasing cell survival.

RSAD2/Viperin is ER/lipid droplet-associated [26], and CMPK2 is mitochondrial [27], positioning them at opposite ends of an ER–mitochondria stress axis. Viperin’s N-terminal amphipathic helix targets the ER/lipid droplets, domains essential for membrane biogenesis and secretory load, suggesting that Viperin may support ER homeostasis under infection induced proteostatic stress [28]. Increased BiP/GRP78 and CHOP, peIF2α elevation, and ATF4 reduction after RSAD2/CMPK2 knockdown indicate UPR remodeling. Although the canonical PERK-eIF2α-ATF4-CHOP cascade often elevates ATF4, contexts exist where sustained eIF2α phosphorylation enforces global translation attenuation with disparate ATF4 dynamics, particularly under combined hypoxia/oxidative stress, pointing to non-canonical UPR tuning in our system. Notably, EBV co-opts UPR effectors, the spliced transcription factor XBP1s directly activates BZLF1, and UPR-inflammasome crosstalk can unlock lytic entry in B-cell lymphomas, providing one route by which organelle stress intersects EBV reactivation [29, 30].

Our transcriptomic analysis revealed a critical role for RSAD2 and CMPK2 in regulating interferon response. EBV reactivation induced a strong interferon response that was severely suppressed by knockdown of either RSAD2 or CMPK2. This attenuation of immune signaling underscores the importance of RSAD2 and CMPK2 in maintaining antiviral defense mechanisms. Interestingly, despite the distinct enzymatic functions of RSAD2 and CMPK2, their knockdowns resulted in highly overlapping transcriptional profiles, suggesting a coordinated regulatory axis. Both genes are highly evolutionarily conserved, occur immediately adjacent to each other and are co-expressed during IFN-signaling. This shows functional cooperation. Both genes modulated a shared set of pathways, including IFN-response, oxidative phosphorylation, electron transport, ATP production, cholesterol biosynthesis, SRP-dependent cotranslational protein targeting to membrane as well as immune-related pathways such as PD1-PD-L1 cancer immunotherapy signaling. The consistent downregulation of genes such as STAT1, and DDX58 (RIG-I) supports the notion that RSAD2 and CMPK2 are upstream regulators of the IFN-signaling cascade. We also observed an increase in Interleukin-1 family signaling after RSAD2 knockdown. These findings point to a dual role for RSAD2 and CMPK2 in immunometabolism, linking antiviral defense to mitochondrial and metabolic reprogramming.

RSAD2 and CMPK2 are enzymes that function together in the generation of the antiviral metabolite ddhCTP. We found that ddhCTP levels were highly elevated by EBV reactivation, and that this was strictly dependent on RSAD2. RSAD2 uses CTP as the substrate to generate ddhCTP, and CTP can be generated by CMPK2, suggesting a direct metabolic handoff from CMPK2 to RSAD2. Interestingly, knockdown of CMPK2 did not reduce ddhCTP levels, suggesting alternative sources of CTP substrate are available for RSAD2. ddhCTP may also be generated from ddhC by other kinases such as UCK2, CMPK1 and NDP to generate ddhCTP [31]. A recent study identified ddhC, as an acute phase reactant in serum of individuals infected with SARS-CoV2 and Influenza A virus [32]. Another study implicated RSAD2 generation of ddhCTP as an inhibitor of NAD-dependent enzymes in the TCA cycle [33], although this finding remains controversial [31]. In our study, we found a strong correlation between EBV reactivation and the RSAD2 dependent generation of ddhCTP.

In addition to ddhCTP, global metabolic profiling revealed many other changes in response to EBV reactivation and knockdown of either CMPK2 or RSAD2. Depletion of CMPK2 led to an increase in inosine/hypoxanthine, a hallmark of purine salvage/catabolism flux which often rises with redox/nucleotide stress [34]. Pathway enrichment for polyamine biosynthesis, Alpha linolenic/linoleic acid metabolism, superoxide degradation, glycolysis- all point towards mitochondrial nucleotide stress and redox changes (mtROS). The rise in nervonic (24:1), lignoceric (24:0), and DHA (22:6) implies elongase/desaturase activity involved in polyunsaturated fatty acid (PUFA) elongation and membrane domain architecture that viruses and IFN programs often modulate. Our data shows a coordinated RSAD2/CMPK2 program that couples mitochondrial nucleotide metabolism and redox to glycolysis and lipid elongation, and IFN-α superimposes a similar metabolic signature (particularly nucleotide and lipid remodeling), indicating the ISG axis is a major driver of these metabolic states.

Consistent with this immune inflammatory theme, our study revealed significant upregulation of both RSAD2 and CMPK2 in SLCLs derived from patients with active MS. These findings support a model in which both genes amplify IFN-driven inflammation while influencing EBV dynamics, potentially exacerbating MS pathogenesis.

Collectively, our study highlights RSAD2 and CMPK2 as central regulators of innate immune reprogramming during EBV infection and reactivation. Their influence extends beyond canonical IFN-signaling to encompass mitochondrial function, ribosome biogenesis, ER-associated unfolded protein response and antiviral metabolites. The interplay between these pathways may be critical for balancing immune activation with cellular homeostasis, and further investigation into their roles could uncover novel therapeutic targets for EBV-associated diseases and autoimmune conditions.

## Materials and Methods

### Human PBMC, Spontaneous Lymphoblastoid Cell Lines and Ethics Statement

Spontaneous lymphoblastoid cell lines (SLCLs) and peripheral blood mononuclear cells (PBMC) from MS patients and controls were obtained from collaborators at the National Institutes of Health (NINDS) and the University of Pennsylvania, Perelman School of Medicine and the Wistar Institute, as described previously [16]. All samples were deidentified and obtained with signed informed consent from participants according to IRB approval process at NIH (89N0045 and CR0045) and UPENN (protocol #816805, IRB #4) and the Wistar Institute (protocol #: 22010335) collaboration agreement.

### Cells and treatments

Mutu I, LCL352, and sLCLs from MS patients were cultured in RPMI supplemented with 10% FBS, 2 mM L-glutamine, 100 U/ml Penicillin and 100 µg/ml Streptomycin. For inducing lytic EBV reactivation, cells were treated with 2 mM sodium butyrate (NaB) and 20 ng/ml TPA (12-O-tetradecanoylphorbol-13-acetate) for 24 h. Mutu I cells were treated with Decitabine, a global hypomethylating agent at final concentration of 7.5 µM for 24, 48 and 72 h. For IFN-A treatment, cells were treated with 10 ng/ml IFN-α2A (STEMCELL Technologies, 78076) for 24 h. For all treatments, either untreated cells or cells treated with DMSO were included as negative control.

### shRNA knockdown of RSAD2 and CMPK2

Lentiviruses were generated by cotransfecting pLKO.1 based shRNA expression plasmids obtained from The Wistar Institute Molecular Screening Facility with plasmids pMD2.G and pSPAX2 in 293T cells using Lipofectamine 2000 transfection reagent (ThermoFisher Scientific). Three days after transfection, supernatant containing the lentivirus was collected, centrifuged to remove cell debris, filtered through a 0.45 micron filter, aliquoted and stored at -80 °C until use. For gene knockdown, cells were resuspended in the lentivirus along with polybrene at a final concentration of 8 µg/ml followed by spinoculation at 450 g at room temperature for 1.5 h. After spinoculation, lentivirus was removed and cells were resuspended in fresh medium. After 48 h, puromycin was added to the culture medium at a final concentration of 2 µg/ml. Fresh medium containing puromycin was added to cells every 2-3 days until collection. For inducing lytic EBV reactivation in cells after knockdown, cells were treated with NaB+ TPA for 24 h as described previously. These cells will be referred to as “Induced” hereafter. Cells not treated with NaB+TPA (uninduced) were also cultured alongside induced cells.

### Gene expression analysis by RT-qPCR

Cells were collected after treatments and washed once with PBS. RNA was isolated from cells using RNeasy plus mini kit. RNA was treated with DNase I (Roche, 4716728001) at room temperature for 20 min to remove residual genomic DNA. RNA was reverse transcribed to generate cDNA using High-capacity cDNA reverse transcription kit (Thermo Fisher, 4368814). RT-qPCR was performed using SYBR green mastermix (Thermo Fisher) in a Quantstudio 7 qPCR machine (Applied Biosystems), and ΔCt method was used for relative quantitation. No-template controls and no-reverse transcriptase controls were included in all RT-qPCR reactions. Data were normalized to the housekeeping gene GUSB. Primer sequences for RT-qPCR are listed in **Table S1**.

### Cell viability

Cell viability was determined by CellTiter-Glo assay (Promega, G7570) as per manufacturer’s instructions. Briefly, cells were lysed using lysis reagent and relative luminescence units (RLU) were measured using CLARIOstar Plus microplate reader (BMG Labtech).

### Western Blot

Cell lysates were prepared in radioimmunoprecipitation assay (RIPA) lysis buffer containing protease and phosphatase inhibitor cocktail (ThermoFisher, 78440). Protein concentration was determined using bicinchoninic acid (BCA) protein assay (Pierce) and lysates were subsequently boiled with 4X Laemmli sample buffer (Bio-Rad) containing β-mercaptoethanol. Proteins were analyzed by SDS-polyacrylamide gel electrophoresis (PAGE) on an 8–16% Tris-glycine precast gel (Invitrogen) and transferred to an Immobilon-P membrane (Millipore). Membranes were blocked in Tris-Buffered Saline containing 5% milk and 0.1% Tween-20, followed by incubation with primary antibody against RSAD2 (Cell Signaling Technology, 13996), CMPK2 (Abcam, 194567), EBNA1 (in-house generated antibody in rabbit), ZTA (in-house generated in rabbit), LMP-1 (Millipore Sigma, MABF2248), EA-D (Millipore Sigma, MAB8186), IRAK1 (Cell Signaling Technology), TRAF6 (Cell Signaling Technology, 8028S), TAK1 (Cell Signaling Technology, 5206), GSDMD (Cell Signaling Technology, 36425T), Caspase 3 (Cell Signaling Technology, 14220), Caspase 1 (Cell Signaling Technology, 3866), eIF2α (Cell Signaling Technology 5324T), p-eIF2α (Cell Signaling Technology 3398S), ATF-4 (Cell Signaling Technology, 11815T) or B-actin (Sigma-Aldrich, A3854). Proteins expressed during ER stress were detected using ER Stress antibody sampler kit (Cell Signaling Technology, 9956T). Membranes were washed with TBST, incubated for 1 h with the goat anti-rabbit IgG-HRP (Biorad, 1706515) or goat anti-mouse IgG-HRP (Biorad, 1706516). Membranes were then washed and detected by enhanced chemiluminescence using Amersham Imager 680 (GE Healthcare).

### Flow cytometry

For flow cytometry analysis, cells were first gated on FSC-A vs. SSC-A to define the primary cell population followed by singlets identification using FSC-A vs. FSC-H (Fig. 4A). All downstream analysis was performed on resulting cell population. Cells were stained using Annexin V-7AAD Apoptosis Detection Kit (ThermoFisher Scientific) as per manufacturer’s instructions and analyzed by FACSymphony A5 SE flow cytometer (BD Biosciences). Data analysis was performed using FlowJo software v10.10.0 (BD Biosciences).

### EBV DNA quantification

Cells were harvested by centrifugation and washed once with PBS. DNA was extracted from cells using DNeasy Blood and Tissue kit (Qiagen, 69504) as per manufacturer’s instructions. DNA concentration was determined using Qubit fluorometer. Quantitative PCR was performed using SYBR green mastermix (ThermoFisher Scientific) using EBNA-1 (EBNA1_F: TCATCATCATCCGGGTCTCC; EBNA1_R: CCT ACAGGGTGGAAAAATGGC); and B-actin (B-actin_F: GCCATGGTTGTGCCATTACA; B-actin_R: GGCCAGGTTCTCTTTTTATTTCTG) primers and relative EBV DNA copies were determined by ΔCt method.

### Chromatin Immunoprecipitation (ChIP)

ChIP was performed to determine EBNA1 binding to EBNA1/CTCF binding site using EBNA1 antibody (generated in-house in rabbits) as described previously [35]. ChIP DNA was assayed by qPCR using primers specific for indicated regions and quantified as % input. Primers for ChIP-qPCR are listed in Table S1.

### RNA-seq

For all the samples, RNA was extracted and treated with DNase I using the DNase treatment kit (Ambion). RNA quality was determined using the Bioanalyzer (Agilent). Only samples with RIN numbers >7.5 were used for further studies. The total RNAseq libraries were made with KAPA Hyper-Prep with rRNA removal by using the Qiagen FastSelect kit, then subjected to 2 × 75-bp high-output sequencing on the Illumina NextSeq 500 platform.\

The read quality was assessed using FASTX (http://hannonlab.cshl.edu/fastx_toolkit/) and FastQC (http://www.bioinformatics.babraham.ac.uk/projects/fastqc) and adapters were trimmed with Cutadapt (https://journal.embnet.org/index.php/embnetjournal/article/view/200). The resulting filtered reads were aligned using Bowtie2 [36] via RSEM V1.3.3 software [37]. Alignment was performed on a reference index created from the Ensemble transcriptome version GRCh37 reference genome for cellular genes and the NCBI Reference Sequence NC_007605.1 for EBV. This enabled estimation of read counts and RPKM values. Subsequently, a comprehensive quality report was generated using MultiQC [38]. Following this, in the read count QC stage, only the genes with ten or more reads in at least one sample were included. Samples were then assessed for outliers using Z-score, Principal Component Analysis (PCA), and correlation plots. Further, raw read counts of all genes were used as input to identify differentially expressed genes between groups using the R package DESeq2 [39]. Gene expression changes were considered significant if they passed the FDR<0.05 threshold. With the significant genes, pathway analysis was performed with QIAGEN’s Ingenuity Pathway Analysis software (IPA, QIAGEN Redwood City, www.qiagen.com/ingenuity) using “Canonical pathways”, “Diseases & Functions”, and “Upstream Regulators” options to identify biological processes and signaling pathways significantly affected by the experimental conditions. Additionally, to obtain a holistic view of pathways associated with the entire gene set, Gene Set Enrichment Analysis (GSEA) [40] was also performed based on Gene Ontology (GO) terms, Molecular Signatures (MSigDB), Hallmark, Reactome, BioCarta, and KEGG pathway databases. Finally, for heatmap visualization of gene expression, we used DESeq2 normalized count values, and other plots were generated using the R package tidyverse (https://joss.theoj.org/papers/10.21105/joss.01686).

### Confocal Imaging

Mutu I cells were treated with sodium butyrate and TPA for 48 h to induce lytic reactivation. Cells were washed with PBS and fixed with 4% (v/v) paraformaldehyde (Thermo Fisher, J19943.K2) for 20 min at room temperature. Cells were then permeabilized in 0.2% Triton X-100 in PBS for 5 min and blocked with PBS containing 10% goat serum for 1 h at room temperature. Primary antibodies for CMPK2 (PA5-34461, ThermoFisher, 1:50) and TOM20 (sc-166755; 1:75) diluted in 1% goat serum were applied to the cells for 1.5 h at room temperature. After 3 washes with PBS containing 0.25% Tween 20, cells were incubated with Alexa 488-, Alexa 568-conjugated secondary antibodies (Invitrogen; 1:800) in 1% goat serum for 1 h at room temperature. Confocal imaging was performed on Leica TCS SP8 X WLL Scanning Confocal Microscope (Leica Microsystems). Images were acquired with 63x,1.4 NA oil immersion objective and the Leica LAS-X software. Confocal z-stacks were acquired with optimal z-intervals according to the Nyquist criterion, covering the whole volume of cells from the basal to the apical region. Images were then compiled by ‘max projection’ before analysis in ImageJ.

### Image Analysis

Image analyses were conducted using ImageJ Fiji (v.2.9.0, National Institutes 598 of Health). Maximum intensity projections were generated from z-stack confocal images. For the measurement of CMPK2 colocalization with TOM20, intensity threshold segmentation was applied to individual cells to identify areas corresponding to CMPK2 and TOM20. Colocalization was assessed using the “JACoP” plugin in ImageJ to calculate Pearson’s coefficient.

### Global metabolite profiling and targeted analysis of ddhCTP, ddhCDP and ddhCMP

RSAD2 and CMPK2 were knocked down in Mutu I cells followed by treatment with sodium butyrate and TPA for 24 h to induce lytic reactivation as described before. Cells were treated with 10 ng/ml IFN-2A for 24 h to induce RSAD2 and CMPK2 expression and were used as positive control. After all treatments, cells were collected and polar metabolites were extracted with ice-cold 80:20 (v/v) methanol/water. For both global and targeted analyses, samples were analyzed by liquid chromatography- mass spectrometry (LC-MS) on a Thermo Scientific Q Exactive Plus mass spectrometer with HESI II probe in-line with a Thermo Scientific Vanquish UHPLC System. Samples were analyzed in a pseudorandomized order. LC separation was performed under HILIC conditions using a ZIC-pHILIC column (150 × 2.1 mm, 5 μm) maintained at 45 °C (EMD Millipore). Mobile phase A was 20 mM ammonium carbonate, 5 µM medronic acid, 0.1% ammonium hydroxide, pH 9.2, and mobile phase B was acetonitrile. Analytical separation was performed at 0.2 ml/min flow rate using the following gradient: 0 min, 85% B; 2 min, 85% B; 17 min, 20% B; 17.1 min, 85% B; and 26 min, 85% B. Specific MS parameters include: sheath gas, 30; auxiliary gas, 5; sweep gas, 0; auxiliary gas heater temperature, 200°C; spray voltage, 3.6 kV for positive/negative polarities; capillary temperature, 325°C; S-lens RF, 65.

For global metabolite profiling, samples were analyzed by either full MS scans with polarity switching (all samples) or full MS/data-dependent MS/MS scans with separate acquisitions for positive and negative polarities (sample pool). Full MS scans were acquired using a scan range of 65 to 975 m/z; 70,000 resolution; automated gain control (AGC) target of 1E6; and maximum injection time (IT) of 100 ms. Data-dependent MS/MS was performed on the 10 most abundant ions; 17,500 resolution; AGC target of 5E4; maximum IT of 50 ms; isolation width of 1.0 m/z; and stepped normalized collision energy of 20, 40, 60. Raw data were processed using Compound Discoverer 3.3 SP3 (Thermo Scientific) with separate analyses for positive and negative polarities. Metabolites were identified by matching accurate mass and retention time to analytical standards or querying MS/MS spectra against the mzCloud spectral database (full match, score > 50; mzCloud.org). Only [M+H]+ and [M-H]- adducts were considered for annotations. Metabolite levels were determined by integrated peak areas using Full MS data. Metabolite levels were corrected for instrument drift using peak areas from technical injections of the sample pool run periodically throughout the analysis and were subsequently normalized to total signal from annotated metabolites in each sample. For targeted analysis of ddhCTP, ddhCDP, and ddhCMP, samples were analyzed by selective ion monitoring (SIM) mode in negative ion mode with an inclusion list (5 ppm mass tolerance) with the theoretical masses for [M-H]- adducts for ddhCTP (463.9667 m/z), ddhCDP (384.0003 m/z), ddhCMP (304.0340 m/z); 70,000 resolution; AGC target of 1E6; maximum IT of 50 ms; and isolation width of 1.0 m/z. Raw data were analyzed using TraceFinder 4.1 (Thermo Scientific). The peak detected for ddhCTP at 11.5 min was confirmed with a reference compound, while the peaks detected for ddhCDP at 10.8 min and ddhCMP at 9.6 min were unconfirmed, since no reference standard was available, but corresponded to the expected elution order of these compounds. Relative quantification was based on integrated peak areas, which were normalized to cell counts for each sample.

### Pathway enrichment analysis for global metabolite profiling

Pathway enrichment analysis for the global metabolite profiling data was performed using the Enrichment Analysis module in MetaboAnalyst 6.0 [41]. Significantly changed metabolites for a given comparison were defined as absolute fold-change greater than 1.5 and Benjamini-Hochberg adjusted p-value (q-value) of less than 0.05. The analysis was performed using the SMPDB metabolite sets, and the reference metabolome corresponded to the set of metabolites annotated in the global metabolite profiling analysis.

## Data Availability

RNAseq dataset is available in the NCBI GEO (GSE). Metabolomics is provided in Supplemental Data.

## Acknowledgements

We would like to thank members of Lieberman lab and the Wistar Institute Cancer Center Shared Resources for Bioinformatics, Genomics, Flow Cytometry and Microscopy Facilities for helpful discussions and excellent assistance with assays and data analysis.

## Funding Information

Funding support was from NIH R01 AI153508-01A1, R01 DE017336, P01 CA281867 (Roberston, PI) to PML, United Sates Department of Defense Award Number: HT9425-23-1-1049 (SSS and PML), AI180133-01 (T. Groves), and The Wistar Institute Cancer Center Support Grant P30 CA010815.

## Notes

### Competing Interest Statement

PML declares competing interest: PML is a founder and advisor to Vironika, LLC, holds patents on EBV inhibitors, and has served as a paid advisory panel member for Sanofi, Merck, GSK, and Pfizer. All other authors declare no competing interest.

